# Loss of catalytic activity and impaired proteostasis in guanosine nucleotide-depleted LRRK2

**DOI:** 10.64898/2026.04.21.719846

**Authors:** Giulia Favetta, Susanne Herbst, Anna Masato, Giulia Tombesi, Lucia Iannotta, Ilaria Battisti, James E. Tomkins, Daniah Trabzuni, Nicoletta Plotegher, Maximiliano G. Gutierrez, Giorgio Arrigoni, Claudia Manzoni, Patrick A. Lewis, Elisa Greggio, Susanna Cogo

## Abstract

Coding mutations in the *Leucine-rich repeat kinase 2* (*LRRK2*) gene represent the most common cause of familial Parkinson’s disease (PD), and are frequently observed in idiopathic PD. In addition, variation around the *LRRK2* locus has been shown to alter PD risk by genome-wide association studies. Disease-causing mutations cluster within the catalytic core of LRRK2 – composed of GTPase (ROC) and serine-threonine kinase domains – and lead to an increase in kinase activity, resulting in hyperphosphorylation of a subset of RAB GTPases and consequent cellular toxicity. However, the interplay between LRRK2 GTPase and kinase domains, and with the surrounding scaffold regions has remained underexplored, with implications for the prediction of on- and off-target effects associated with kinase inhibition. To address this gap, here we dissected the contributions of kinase, GTPase and scaffold domains to LRRK2 function in murine macrophages and tissues expressing endogenous levels of GTP/GDP-binding deficient Lrrk2 T1348N. Guanosine nucleotide-free Lrrk2 is devoid of both GTPase and kinase activities but maintains the scaffold shell, leading to significant reshaping of Lrrk2 interactome and engagement in novel interactions. This altered functional state leads to impaired autophagy and accumulation of enlarged lysosomes and autophagic cargo in macrophages and kidneys. Since pharmacological inhibition of LRRK2 is under clinical evaluation, our results reveal retained scaffold functions upon loss of catalytic activity that warrant careful consideration.

## Introduction

Parkinson’s disease (PD) is a progressive neurodegenerative disorder characterized by prominent disruption of movement as well as non-motor symptoms^1^. The clinical manifestations of PD are accompanied by neuronal loss in the central nervous system (CNS), predominantly in the *substantia nigra pars compacta*, and the accumulation of intracellular inclusions made up of protein aggregates and cellular membrane components, known as Lewy bodies^2,3^. The etiology and pathogenesis of PD are complex, however the association of genetic variants with enhanced risk of disease has provided important insights into the cellular events leading to neuronal cell death^4^. A key contributor to genetic risk for PD is the *LRRK2* gene located on chromosome 12, encoding the multidomain signaling protein Leucine Rich Repeat Kinase 2 (LRRK2)^5^. *LRRK2* is a pleiotropic locus for PD, with coding mutations linked to autosomal dominant PD^6,7^, and common non-coding variation associated with increased risk of disease in the general population^8,9^. The cellular function of LRRK2, and how this goes awry during disease, is therefore of great interest – especially in the light of ongoing clinical trials targeting LRRK2^10^.

Although PD primarily affects the CNS, LRRK2 is prominently expressed in the myeloid lineage, including not only CNS-resident microglia, but also peripheral macrophages^11–13^. Hence, macrophage systems are a suitable model to interrogate LRRK2-dependent mechanisms. Data from a range of cellular and *in vivo* systems suggest that LRRK2 has an important role in the response to lysosomal damage and the regulation of macroautophagy^14–17^, and LRRK2 has been implicated in additional cellular processes including mitochondrial function^18^, cytoskeletal dynamics^19^ and synaptic activity^20,21^. Connecting these apparently disparate functions is the ability of LRRK2 to regulate membrane trafficking via direct phosphorylation of a subset of RAB GTPases, including RAB8A, RAB10, and RAB12 (among others)^22,23^. In addition, RAB29 and RAB32 have been implicated as upstream recruiters/activators that control LRRK2 membrane localization and activity^24–26^. Importantly, coding mutations in LRRK2 associated with PD result in increased phosphorylation of RAB substrates^27^.

LRRK2 is one of three proteins within the human proteome that possesses both kinase and GTPase (Ras of Complex, ROC) domains^28^, with these two activities being tightly connected^29^. Artificial mutations within the GTP binding pocket of LRRK2 (e.g., a lysine to alanine substitution at position 1347, and a threonine to asparagine substitution in position 1348) that block the ability of LRRK2 to bind guanosine nucleotides, have previously been used to investigate how disruption of the GTPase activity influences LRRK2 kinase activity and function^30–33^. To further clarify how guanosine nucleotide binding regulates LRRK2 physiological function, here we investigated the biochemical and cellular consequences of the T1348N variant at an endogenous level in RAW 264.7 macrophages, primary macrophages, and tissues from Lrrk2 T1348N knock-in (KI) mice. We uncovered that nucleotide-binding-deficient LRRK2, while both GTPase and kinase inactive, retains scaffold properties that are sufficient to reshape protein interactions and perturb autophagy-lysosome pathways in macrophages and kidneys, suggesting that it is not functionally equivalent to a complete loss of LRRK2 function. This work is particularly relevant in light of the presence of a naturally occurring variant at residue 1348 in human populations (T1348P), which is predicted to impair the ability of LRRK2 to bind GTP^33^, and in the context of LRRK2-targeted drug discovery, where clinical trials focused on novel LRRK2 kinase inhibitors are highlighting the importance of a comprehensive understanding of LRRK2 biology beyond its kinase activity^10,34^.

## Results

### Loss of guanosine nucleotide binding lowers Lrrk2 levels and impairs canonical Lrrk2-Rab signaling in macrophages

To determine how impaired guanosine nucleotide binding within the ROC GTPase domain affects LRRK2 steady-state levels and kinase activity, we examined murine RAW 264.7 macrophage cells harboring the T1348N mutation in endogenous Lrrk2, and primary bone marrow-derived macrophages (BMDMs) isolated from Lrrk2 T1348N KI mice. T1348N is a P-loop substitution engineered to disrupt GTP/GDP binding and ROC GTPase activity^32,33,35^ (Fig. 1A-A’). Consistent with previous studies using ectopically expressed proteins^30,32,36^, Lrrk2 T1348N steady-state protein abundance was markedly reduced. In RAW 264.7 cells, Lrrk2 T1348N levels were decreased by approximately 80% as compared to wild-type (WT), while remaining detectable above the knockout (KO) control cell line, included as a reference for Lrrk2 loss of function (Fig. 1B-B’). A comparable reduction was observed in BMDMs (Fig. 1C-C’), indicating that decreased steady-state abundance is conserved across macrophage systems. We next assessed Lrrk2-mediated phosphorylation of Rab10 at Thr73 (pRab10 T73), a canonical LRRK2 substrate site. Treatment with lysosomotropic agents, including L-leucyl-L-leucine methyl ester (LLOMe) and chloroquine (CQ), significantly enhances lysosomal recruitment of LRRK2 and LRRK2-dependent RAB10 T73 phosphorylation, initiating lysosomal repair mechanisms^15,16,23,37,38^. In both RAW 264.7 cells and BMDMs, pRab10 T73 signal was not detected in the Lrrk2 T1348N cells relative to WT in unstimulated conditions, nor could it be induced by treatment with CQ, as shown in Fig. 1D-E. Importantly, Rab10 T73 phosphorylation in WT RAW 264.7 cells was abolished by treatment with the LRRK2 kinase inhibitor MLi-2^39^ (Fig. 1D). We additionally tested the ability of LLOMe to induce Lrrk2, Rab8A and Rab10 recruitment to the lysosomal compartment by monitoring puncta formation via immunofluorescence. While Lrrk2 shows a diffuse cytoplasmic pattern of expression in unstimulated conditions, when treated with LLOMe WT cells displayed robust Lrrk2, Rab8A and Rab10 puncta formation, indicative of Lrrk2-dependent recruitment to the lysosomal compartment, whereas T1348N and KO cells failed to show Lrrk2 and Rab8A/Rab10 recruitment (Fig. 1F-I). Together, these data are consistent with the loss of Lrrk2-dependent Rab phosphorylation and suggest that the GTP/GDP binding-defective T1348N mutant is catalytically inactive and unable to initiate canonical LRRK2 signaling. However, the differential protein levels between WT and T1348N Lrrk2 preclude resolving whether these effects are predominantly due to the reduction in absolute protein levels, the nucleotide-binding disruption within ROC, or a combination of both.

**Figure 1.**
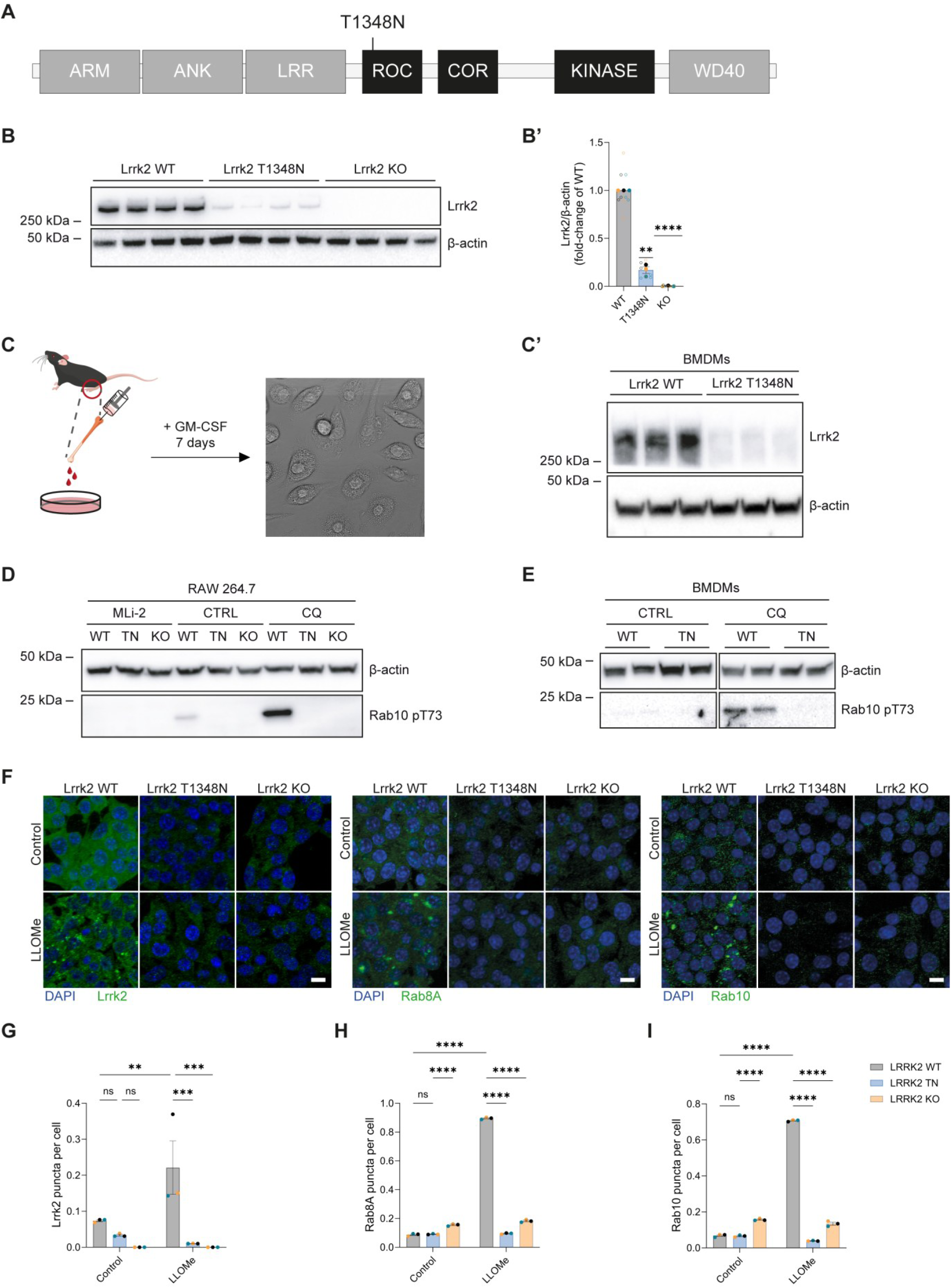
The ROC P-loop mutation T1348N reduces Lrrk2 abundance and impairs Rab phosphorylation and recruitment to damaged lysosomes in macrophages. (**A**) Schematic of Lrrk2 domain organization indicating the position of the ROC P-loop substitution T1348N. The catalytic core is shown in black; scaffold domains are indicated in grey. (**B-B’**) Immunoblots of endogenous Lrrk2 steady-state levels in murine RAW 264.7 macrophage cell lines (WT, Lrrk2 T1348N, and Lrrk2 KO), with β-actin as loading control (**B**), and corresponding quantification of Lrrk2 abundance normalized to WT within each independent experiment (*n* = 3, **B’**). (**C-C’**) Schematic of the protocol for differentiation of primary bone marrow-derived macrophages (BMDMs) from WT and Lrrk2 T1348N KI mice using GM-CSF (granulocyte-macrophage colony-stimulating factor; 50 ng/mL) for 7 days (representative brightfield image for illustration purposes only, **C**), and immunoblots of endogenous Lrrk2 steady-state levels in BMDMs, with β-actin as loading control (**C’**). (**D**-**E**) Immunoblots of Lrrk2-mediated phosphorylation of Rab10 at Thr73 (pRab10 T73) in RAW 264.7 cells (**D**) treated with vehicle (CTRL), chloroquine (CQ, 50 μM, 2 h) or the Lrrk2 inhibitor MLi-2 (0.1 μM, 90 min), and BMDMs (**E**) under vehicle (CTRL) or CQ (50 μM, 2 h). β-actin is used as loading control. (**F-I**) Representative immunofluorescence images of RAW 264.7 WT, Lrrk2 T1348N and Lrrk2 KO cells treated with vehicle or L-leucyl-L-leucine methyl ester (LLOMe; 1 mM, 30 min), showing stimulus-dependent puncta formation/recruitment of Lrrk2 (left), Rab8A (middle) and Rab10 (right) (**F**). Scale bars 10 μm. Quantification shows the number of Lrrk2-positive (**G**), Rab8A-positive (**H**) and Rab10-positive (**I**) puncta per cell under vehicle and LLOMe conditions. For each biological replicate (*n* = 3), multiple fields were quantified and averaged to yield one value per replicate. Data are presented as mean ± SEM unless otherwise stated; solid dots represent independent biological replicates (mean of technical replicates, shown as open dots, within each experiment). Statistical significance: ** p < 0.01, *** p < 0.001, **** p < 0.0001. Statistical test: (**B’**) two-tailed one-sample t test. (**G-I**) two-way ANOVA with Tukey’s multiple comparisons.

### Reduced Lrrk2 T1348N levels are not explained by altered turnover or solubility

The significant reduction in Lrrk2 T1348N protein levels prompted us to ask whether differences in transcript levels, protein turnover, or solubility could account for its decreased steady-state abundance. Lrrk2 T1348N mRNA levels were reduced by 25% in RAW 264.7 cells compared to WT (Fig. 2A), a change that may contribute to the more pronounced (approximately 80%) decrease observed at the protein level, consistent with the often-nonlinear relationship between mRNA abundance and protein output^40,41^. Alternatively, additional post-transcriptional mechanisms could account for the observed reduction in protein abundance. Therefore, we next assessed whether increased protein turnover could underlie the T1348N deficit. Cycloheximide (CHX) chase experiments revealed stable Lrrk2 levels over a 48-hour window in both WT and T1348N Lrrk2 cell lines (Fig. 2B-B’). Previous studies in Tet-on HEK293 cells also suggested a long, apparent half-life of the protein of approximately 38 hours^42^, although cell-type specificity might exist. Because similar net turnover does not exclude preferential engagement of specific proteolytic routes, we next tested whether blocking major degradation pathways would selectively increase T1348N levels. Pharmacological inhibition of proteasomal (MG132) or lysosomal (CQ) degradation did not selectively rescue Lrrk2 T1348N levels within the tested timeframe (Fig. 2C-C’), suggesting enhanced clearance through either pathway is not the primary driver of reduced steady-state levels. Consistent with this, WT and T1348N Lrrk2 displayed comparable solubility (Fig. 2D-D’; independent validation by Dr Francesca Pischedda and Dr Giovanni Piccoli, University of Trento), arguing against a major shift toward insoluble/aggregated species within the detected protein pool. In addition, using a cellular thermal shift assay (CETSA) (Fig. S1), we did not observe consistent changes in the denaturation profiles of WT vs T1348N Lrrk2, although CETSA readouts can also reflect changes in binding partners/complex assembly rather than intrinsic folding stability^43^. Together, these analyses indicate that the marked reduction of Lrrk2 T1348N at steady state is not caused by accelerated bulk turnover, preferential proteasomal/lysosomal clearance, or major changes in apparent thermal stability or solubility, but could be, at least partially, explained by lower transcript expression.

**Figure 2.**
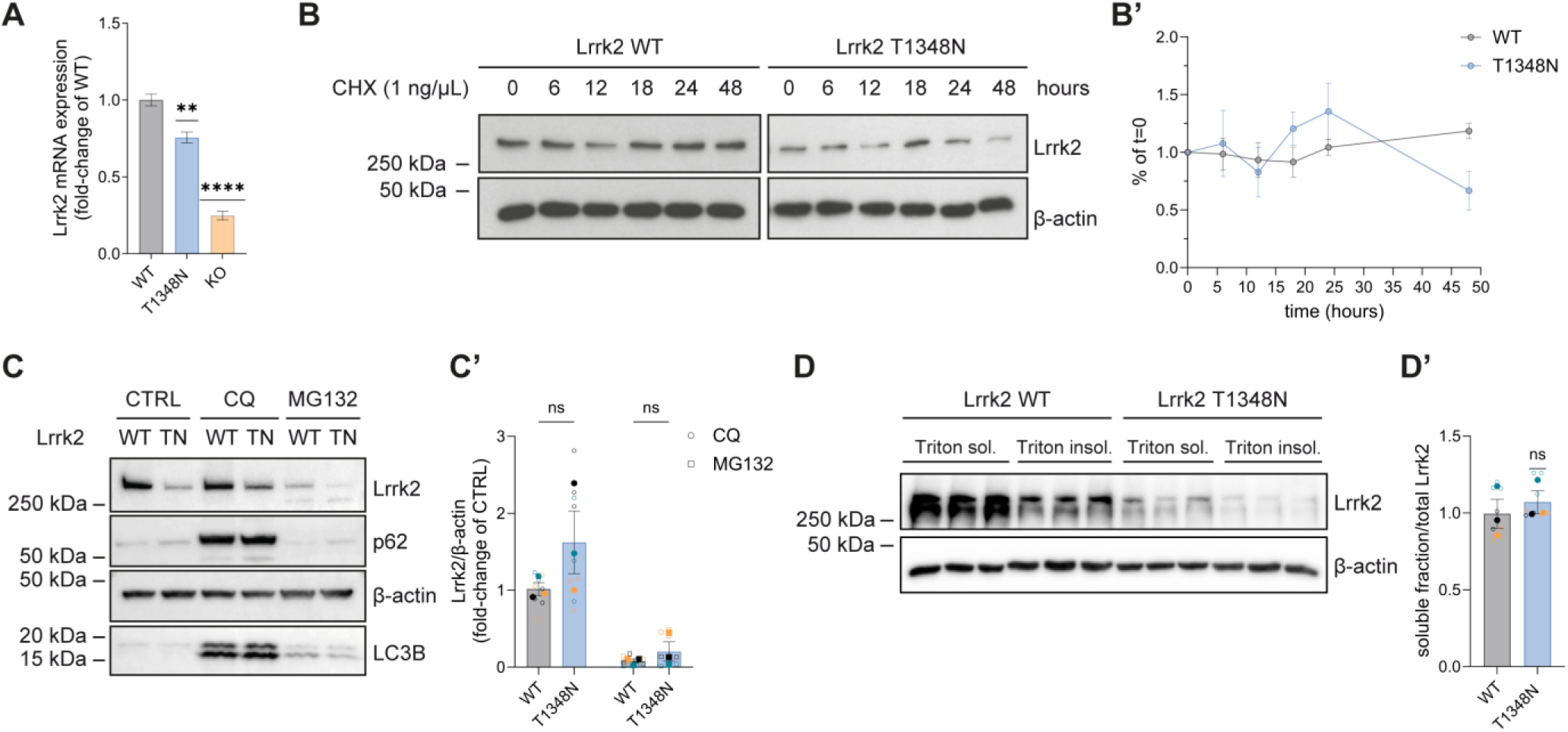
Reduced Lrrk2 T1348N abundance is not explained by altered turnover or solubility. (**A**) RT-qPCR analysis of Lrrk2 transcript levels in murine RAW 264.7 cells (WT, Lrrk2 T1348N, and Lrrk2 KO), normalized and expressed relative to WT (*n* = 5 replicates). (**B-B’**) Cycloheximide (CHX) chase in RAW 264.7 WT and Lrrk2 T1348N cells (CHX, 1 ng/μL; time points as indicated), with immunoblot detection of Lrrk2 and β-actin as loading control (**B**). Densitometric quantification shows Lrrk2/β-actin expressed as a fraction of the t = 0 signal within each genotype (**B’**). (**C-C’**) Immunoblots of Lrrk2 levels in RAW 264.7 WT and Lrrk2 T1348N cells following inhibition of major degradation pathways (MG132, 20 μM, 16 h for proteasome inhibition; CQ, 50 μM, 16 h for lysosomal neutralization), with β-actin as loading control (**C**). p62 and LC3B are shown as pathway response markers. Quantification of Lrrk2 abundance across treatments is expressed as fold-change relative to vehicle (CTRL) within each genotype (*n* = 3 independent experiments, **C’**). (**D-D’**) Solubility fractionation of WT and T1348N Lrrk2, assessing distribution between Triton X-100 -soluble and -insoluble fractions (**D**); quantification shows soluble Lrrk2 over total, normalized to WT within each independent experiment (*n* = 3, **D’**). Data are presented as mean ± SEM; solid dots represent independent biological replicates (mean of technical replicates, shown as open dots, within each experiment). Statistical significance: ns, not significant; ** p < 0.01, **** p < 0.0001. Statistical tests: (**A**, **D’**) one-sample t test; (**B’**) two-way ANOVA with Tukey’s multiple comparisons; (**C’**) two-way ANOVA with Šídák’s multiple comparisons.

### LRRK2 T1348N disrupts 14-3-3 interactions while increasing binding with chaperones

We next investigated whether impaired guanosine nucleotide binding rewires LRRK2-associated protein interaction networks in modes that could provide insight into its reduced abundance and potential functional consequences. To compare WT and T1348N LRRK2 interactomes, we performed affinity purification-mass spectrometry (AP-MS) of 3xFlag-tagged WT and T1348N LRRK2 expressed in HEK-293T cells (Fig. 3A). Importantly, WT and T1348N LRRK2 were expressed at matched levels and recovered at comparable amounts in anti-Flag immunoprecipitation, minimizing confounding effects of unequal bait abundance. Differential abundance analysis revealed marked remodeling of the LRRK2 interactome upon loss of guanosine nucleotide binding (Fig. 3B and Supplementary Data 1). Among the most prominent changes, six 14-3-3 proteins were underrepresented in LRRK2 T1348N complexes, whereas molecular chaperones and chaperone-associated proteins, including members of the HSP70 family, were enriched or in greater abundance with the mutant protein (Fig. 3B). These findings are consistent with reduced phosphorylation at LRRK2 Ser910 and Ser935, the primary docking sites for 14-3-3 proteins^44–47^, and suggest that the T1348N variant engages protein quality-control pathways more strongly than WT LRRK2. This is in line with previous work implicating chaperone pathways in LRRK2 proteostasis, including protein folding and stability control^48–50^. Of note, cytoskeletal protein interactors were not depleted in T1348N LRRK2, in fact the direction of effect was toward enrichment in the mutant form, including for proteins associated with the actin cytoskeleton, such as DBN1, a known LRRK2 interactor^51^. Ribosomal proteins showed a more heterogeneous pattern of differential abundance, although none of these interactors reached statistical significance (Fig. S2). In parallel, analysis of genotype-biased protein detection highlighted a subset of interactors preferentially recovered with either WT or T1348N LRRK2 (Fig. 3C). Notably, the E3 ubiquitin ligase TRIM32 (Tripartite Motif-Containing Protein 32) was reproducibly detected among T1348N-exclusive interactors, suggesting that loss of nucleotide binding may expose or favor interactions with proteostasis- and autophagy-related proteins. To test whether stimuli relevant to LRRK2 function further modulate this rewired interaction network, we validated selected interaction changes by anti-Flag immunoprecipitation followed by immunoblotting, in the absence or presence of CQ treatment to perturb lysosomal homeostasis, and nutrient starvation to enhance autophagic flux (Fig. 3D-D’). This confirmed reduced 14-3-3 binding and increased association with chaperone proteins in LRRK2 T1348N complexes. Moreover, association of TRIM32 with LRRK2 T1348N was increased upon CQ treatment and reduced under starvation (Fig. 3D-D’). Given that TRIM32 has been reported to regulate autophagy through ULK1/AMBRA1-linked mechanisms^52–54^, these data suggest that the T1348N variant may engage proteostatic and autophagy-related mechanisms distinct from those of WT LRRK2.

**Figure 3.**
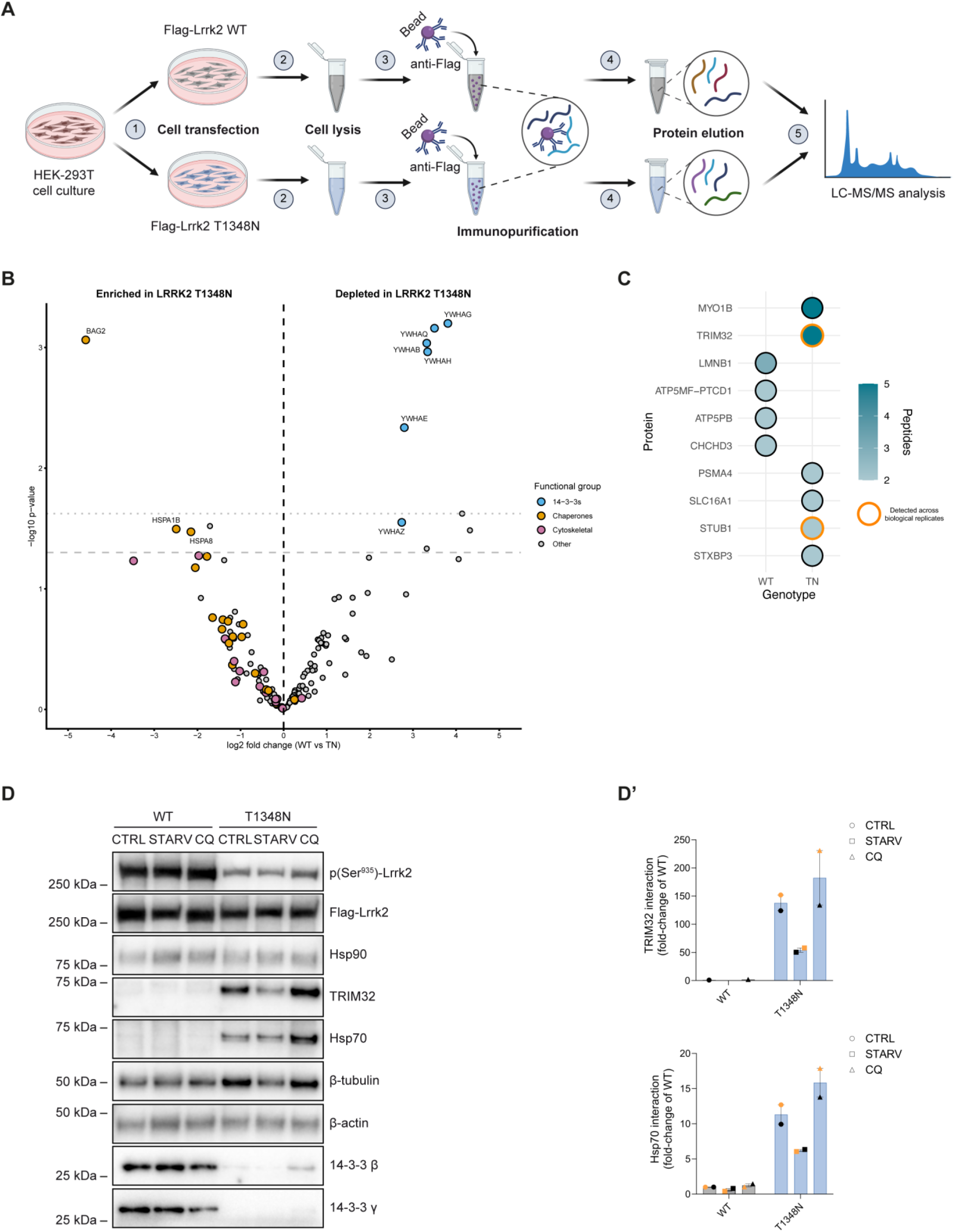
LRRK2 T1348N disrupts 14-3-3 binding and enhances associations with chaperones and autophagy regulators. (**A**) Schematic overview of the AP-MS workflow: anti-Flag affinity purification of 3xFlag-LRRK2 WT or T1348N expressed in HEK-293T cells, followed by LC-MS/MS analysis. (**B**) Volcano plot showing differential enrichment of co-purified proteins with 3xFlag-LRRK2 WT or T1348N under basal conditions. Data are plotted as log2 fold-change (WT/T1348N) against -log10 p-value from *n* = 2 biological replicates per genotype; negative fold changes indicate enrichment with LRRK2 T1348N, whereas positive values indicate reduced association with LRRK2 T1348N. 14-3-3, chaperone and cytoskeletal proteins are highlighted, and labelled when nominal p<0.05 indicated by dashed horizontal line. Adjusted p<0.05 threshold indicated with dotted horizontal line. (**C**) Dot plot showing genotype-exclusive proteins detected above QC threshold in either LRRK2 WT or T1348N AP-MS datasets. Dot fill indicates peptide count, and orange outline indicates proteins detected across biological replicates (*n* = 2), as indicated in the legend. Reproducible T1348N-exclusive proteins include TRIM32 and STUB1. (**D-D’**) Immunoblot validation of selected AP-MS hits in anti-Flag immunoprecipitates from HEK-293T cells expressing 3xFlag-LRRK2 WT or T1348N under control (CTRL), starvation (STARV, 16 h serum deprivation followed by 2 h in HBSS), or CQ treatment (50 μM, 16 h), including assessment of LRRK2 phosphorylation at Ser935, chaperones (Hsp90/Hsp70), TRIM32, and 14-3-3 proteins (γ and β) (**D**). Quantification of TRIM32 and Hsp70 co-purification, expressed as fold-change relative to WT within each condition (*n* = 2 technical replicates, **D’**). Each dot represents one quantified interactor in AP-MS plots or one technical replicate in immunoblot quantifications. Data are presented as mean ± SEM where quantified.

### Lrrk2 T1348N impairs autophagy and drives age-dependent accumulation of the autophagic cargo p62 in kidneys

Given the enrichment of TRIM32 in the interactome analysis, we next tested whether the Lrrk2 T1348N variant alters autophagy-lysosome pathway readouts in cellular and mouse models. Because the autophagy receptor SQSTM1/p62 is normally degraded together with its cargo, its accumulation is commonly used as a proxy for reduced autophagic clearance/turnover^55,56^, although transcriptional upregulation can also contribute to increased p62 levels^57^. In Lrrk2 T1348N RAW264.7 cells, p62 levels were significantly elevated compared with WT and Lrrk2 KO at basal conditions (Fig. 4A-A’), and this increase was recapitulated in primary BMDMs isolated from Lrrk2 T1348N KI mice (Fig. 4B-B’). To test whether altered turnover could be the cause of p62 accumulation, we monitored the autophagic flux in RAW 264.7 cells by following the maturation of the autophagosome marker LC3B (Microtubule-associated protein 1A/1B-light chain 3B), using the tandem fluorescent mCherry-GFP-LC3 reporter. In this system, GFP is quenched upon delivery to acidic autolysosomes while mCherry remains stable, allowing LC3-positive structures to be stratified according to their acidification status^58–60^. To probe flux, we treated RAW 264.7 cells with CQ, which elevates lysosomal pH and inhibits lysosome-dependent degradation, thereby increasing the fraction of GFP-retaining puncta in flux-competent cells^61^. Accordingly, CQ increased the fraction of mCherry-GFP-positive/mCherry-positive LC3B puncta in WT and Lrrk2 KO cells, whereas Lrrk2 T1348N cells showed no reporter shift despite starting from a basal level comparable to WT (Fig. 4C). Importantly, CQ induced the expected accumulation of autophagy markers, i.e., p62 and LC3B, in immunoblot assays (Fig. 2C), indicating effective pathway engagement. This suggests that the attenuated reporter response in T1348N cells likely reflects altered lysosome-dependent processing/acidification of LC3B-positive structures. Together with p62 accumulation (Fig. 4A-A’, 4B-B’), this profile is most compatible with a perturbation at or downstream of the autolysosomal step, such as impaired lysosomal maturation/acidification and/or reduced lysosomal degradative capacity, rather than reduced autophagosome biogenesis^62,63^. We next asked whether the increase in p62 was conserved *in vivo* and varied with age by analyzing p62 levels in brain, kidneys and lungs from WT and Lrrk2 T1348N KI mice at 1, 6 and 12 months. We sought to compare these tissues since high Lrrk2 expression and mutation- as well as tissue-specific phenotypes have been previously reported in Lrrk2 genetic models^64–67^. In pooled lysates, p62 levels showed no consistent genotype-dependent change in brain and lungs, whereas kidney lysates from T1348N mice exhibited higher p62 levels at all time points (Fig. 4D). Further analysis of individual kidney samples confirmed an age-dependent increase in p62 in T1348N mice at 6 and 12 months (Fig. 4E-E’’, 4G). Notably, Lrrk2 protein levels were reduced across tissues (Fig. 4D-F), consistent with the decreased steady-state abundance observed in macrophage models (Fig. 1). Altogether, we found that, despite the loss of enzymatic activities, Lrrk2 T1348N is associated with autophagy-lysosome perturbations that diverge from the Lrrk2 KO phenotype, as p62 accumulation and impaired autophagic flux are observed in Lrrk2 T1348N but not Lrrk2 KO RAW 264.7 macrophages (Fig. 4A-A’ and 4C). The increase in p62 levels is recapitulated in primary BMDMs (Fig. 4B-B’), and Lrrk2 T1348N KI mice develop an age-dependent kidney p62 phenotype (Fig. 4E-E’’, 4G). Collectively, these data challenge a simple loss-of-function interpretation of the T1348N mutant, consistent with residual effects exerted by the non-enzymatic (interaction/scaffold-like) domains.

**Figure 4.**
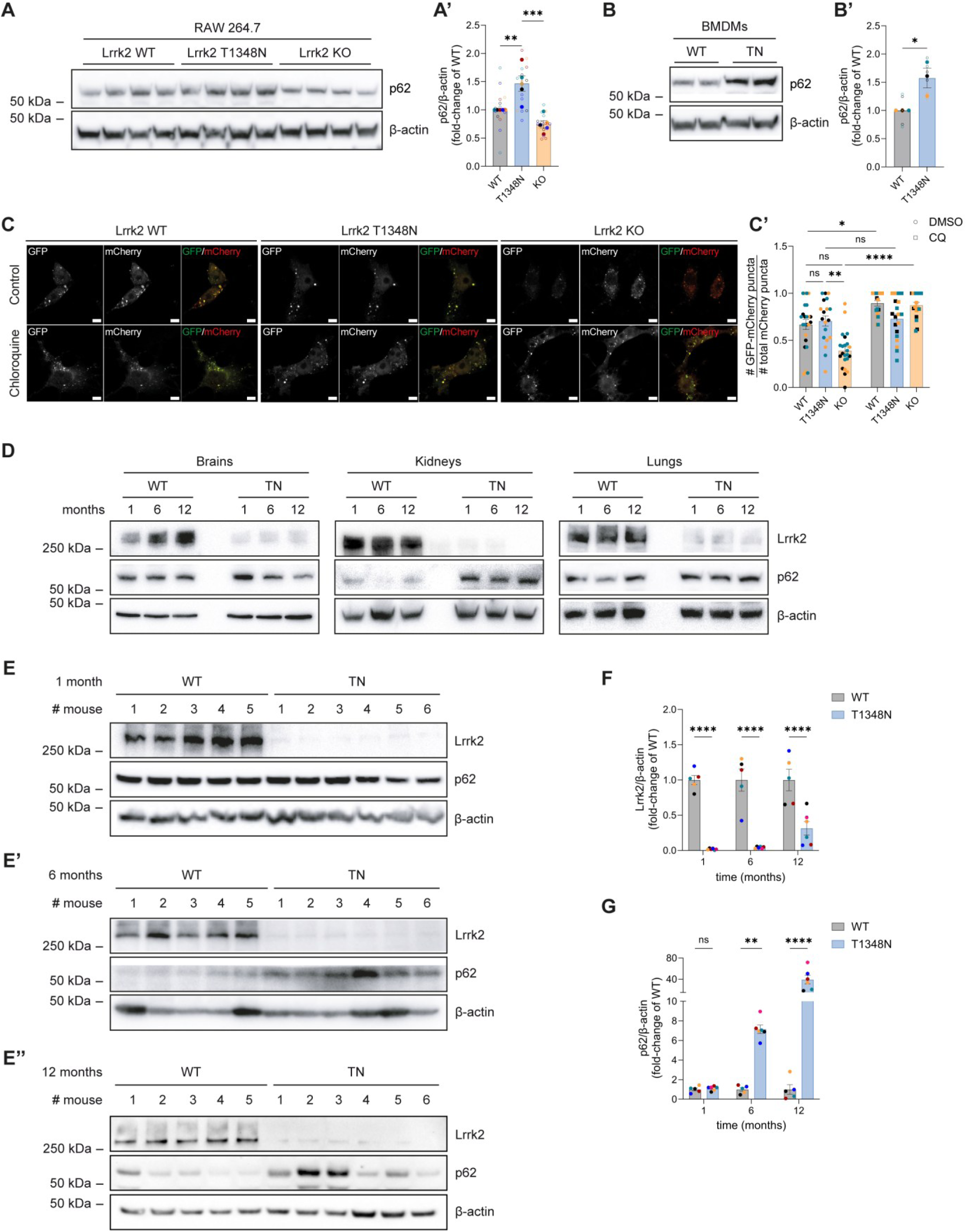
Lrrk2 T1348N perturbs autophagy-lysosome homeostasis and drives age-dependent p62 accumulation in kidneys. (**A-A’**) Immunoblots of p62 levels in RAW 264.7 macrophage cell lines (WT, Lrrk2 T1348N, and Lrrk2 KO), with β-actin as loading control (**A**), and corresponding quantification relative to WT (*n* = 5 independent experiments, **A’**). (**B-B’**) Immunoblots of p62 levels in primary BMDMs from WT and Lrrk2 T1348N KI mice (**B**), with quantification relative to WT (*n* = 3 independent experiments, **B’**). (**C-C’**) Autophagic flux assessment in RAW 264.7 WT, Lrrk2 T1348N, and Lrrk2 KO cells expressing the tandem mCherry-GFP-LC3 reporter under vehicle or CQ treatment (50 µM, 16 h). Representative images show GFP, mCherry, and merged channels (**C**); quantification shows the fraction of GFP-mCherry-positive puncta normalized to total mCherry puncta per cell (*n* = 3 independent experiments, **C’**). (**D**) Immunoblots of Lrrk2 and p62 levels in brain, lung, and kidney lysates from WT and Lrrk2 T1348N KI mice collected at 1, 6, and 12 months (pooled samples per condition), with β-actin as loading control. (**E-E’’**) Immunoblots of individual kidney lysates from WT and T1348N mice at 1 month (**E**), 6 months (**E’**), and 12 months, with β-actin as loading control (**E’’**). (**F-G**) Densitometric quantification of Lrrk2 (**F**) and p62 (**G**) levels in kidneys from (**E-E’’**), expressed relative to WT at each age (*n* = 5 WT mice, *n* = 6 Lrrk2 T1348N mice). Data are presented as mean ± SEM; solid dots represent independent biological replicates. Statistical significance: ns, not significant; * p < 0.05, ** p < 0.01, *** p < 0.001, **** p < 0.0001. Statistical tests: (**A’**) one-way ANOVA with Tukey’s multiple comparisons; (**B’**) unpaired two-tailed Student’s t test; (**C’**) Kruskal-Wallis with Dunn’s multiple comparisons; (**F**, **G**) two-way ANOVA with Šídák’s multiple comparisons.

### Female Lrrk2 T1348N mice show Trim32 and p62 accumulation in kidneys, accompanied by lysosomal enlargement

To further characterize the kidney-selective phenotype observed in Lrrk2 T1348N KI mice (Fig. 4D-E), and to determine whether the autophagy-related interactor TRIM32 emerged from the AP-MS dataset is also altered *in vivo* (Fig. 3), we analyzed kidneys from 12-month-old animals. Immunoblotting revealed increased Trim32 and p62 abundance in Lrrk2 T1348N kidney lysates compared to WT (Fig. 5A-B), consistent with altered autophagy-lysosome homeostasis in this tissue. Notably, within the T1348N cohort, p62 accumulation was more pronounced in females than in males (Fig. 5C’), indicating a sex bias in the magnitude of the phenotype. To understand the spatial distribution of these changes, we performed immunofluorescence on kidney sections from female WT and Lrrk2 T1348N mice (Fig. 5D) and quantified the abundance of p62, Trim32 and the endolysosomal marker Lamp1 (Lysosome-associated membrane protein 1) by integrated density. Compared to WT, Lrrk2 T1348N kidneys exhibited higher p62, Trim32 and Lamp1 levels, accompanied by enlargement of Lamp1-positive vesicles and increased p62-Lamp1 co-localization (Fig. 5E-H), compatible with a stalled autophagic flux. Taken together, these findings link the T1348N genotype to a kidney-specific defect in lysosome-dependent handling of autophagic cargo, accompanied by Trim32 upregulation, and a female-biased increase in p62, consistent with late-stage macroautophagy dysfunction *in vivo*^63^.

**Figure 5.**
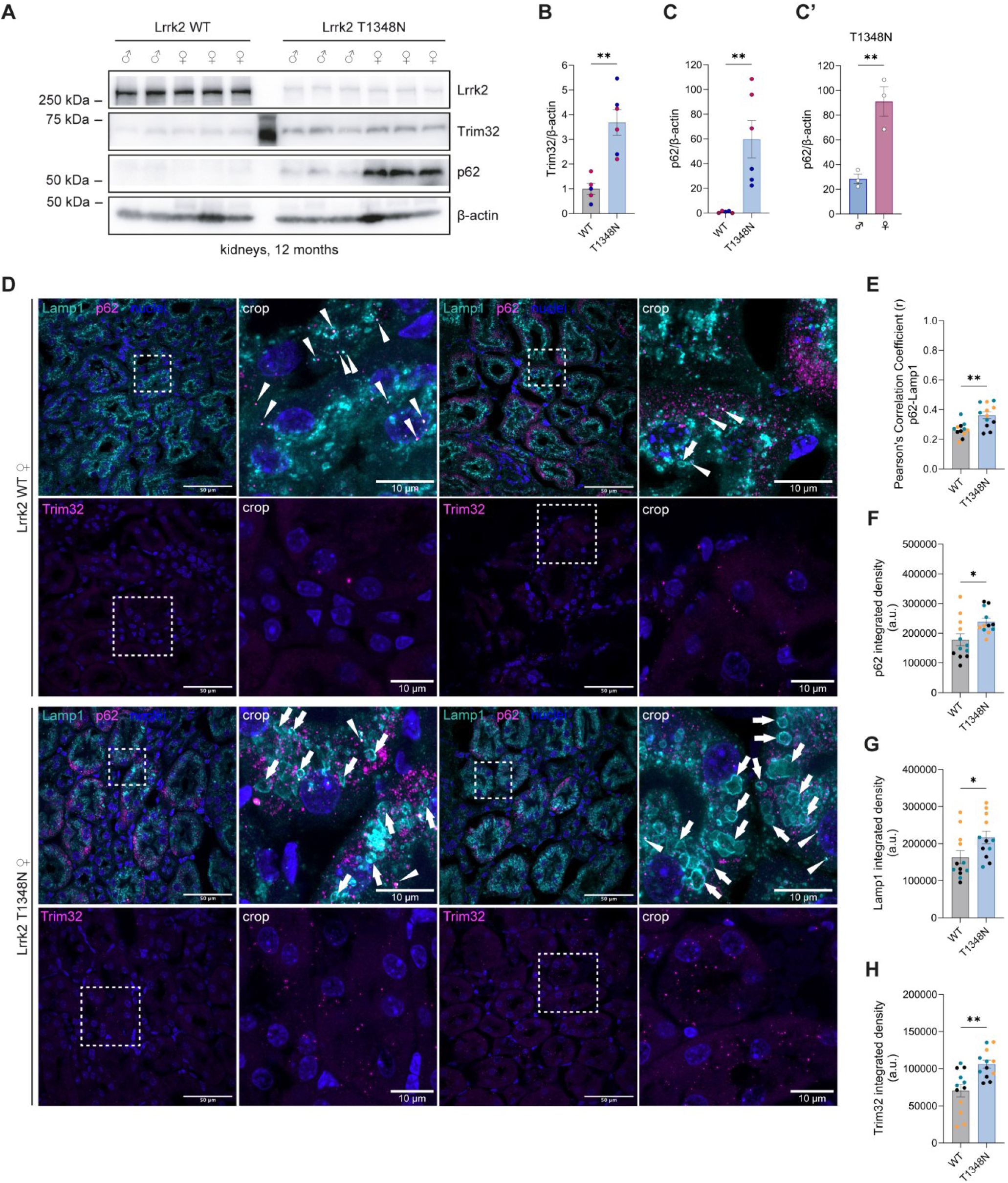
Trim32 upregulation accompanies lysosomal enlargement and sex-biased p62 accumulation in T1348N kidneys. (**A**) Representative immunoblots of Lrrk2, Trim32, and p62 levels in kidney lysates from 12-month-old WT and T1348N mice, with β-actin as loading control. (**B-C’**) Quantification of Trim32 (**B**) and p62 (**C**) normalized to β-actin and expressed relative to WT (*n* = 5 WT mice, *n* = 6 Lrrk2 T1348N mice). p62 levels are also shown stratified by sex within the T1348N cohort (female *n* = 3 vs male *n* = 3, **C’**). (**D**) Representative immunofluorescence images of kidney sections from female WT and T1348N mice stained for Lamp1, p62, and Trim32, with nuclei counterstain. Boxed regions are shown as magnified insets. Scale bars 50 µm (main images) or 10 µm (crops), as indicated. (**E**) Quantification of p62-Lamp1 overlap (Pearson’s correlation coefficient) in kidney sections. (**F-H**) Quantification of p62 (**F**), Lamp1 (**G**), and Trim32 (**H**) signals by integrated density in kidney sections. Data are presented as mean ± SEM; dots represent individual mice. For immunofluorescence quantifications (**E**-**H**), 4 fields per mouse were quantified and averaged to generate one value per mouse (*n* = 3 WT mice, *n* = 3 T1348N mice; see STAR Methods). Statistical significance: * p < 0.05; ** p < 0.01. Statistical tests: (**B-C’**, **E-H**) unpaired two-tailed Student’s t test.

## Discussion

In this study, we characterized the consequences of genetic disruption of guanosine nucleotide binding on the biochemical and cellular functions of LRRK2, using complementary physiological models including Lrrk2 T1348N KI RAW 264.7 macrophages, alongside primary BMDMs and tissues from Lrrk2 T1348N KI mice. By focusing on endogenous expression systems, our work provides mechanistic insight into how loss of GTP/GDP binding reshapes LRRK2 regulation and downstream pathways under physiological conditions. Collectively, we show that the T1348N substitution reduces Lrrk2 steady-state abundance, abrogates canonical Rab phosphorylation and lysosomal recruitment, remodels the LRRK2 interactome, and is associated with tissue- and sex-dependent autophagy-lysosome phenotypes that are distinct from those observed with Lrrk2 KO.

Consistent with previous observations in cellular overexpression systems and Lrrk2 T1348N KI mouse embryonic fibroblasts (MEFs)^30,32,68^, loss of nucleotide binding results in a marked reduction in steady-state Lrrk2 protein levels in both mouse cells and tissues (Fig.1, Fig.4E-G). This decrease, however, cannot be explained by enhanced protein degradation, altered turnover or changes in solubility (Fig. 2B-D’). Instead, the decrease in protein abundance correlates with a reduction in Lrrk2 mRNA levels (Fig. 2A). The T1348N substitution lies within an internal coding exon of Lrrk2 and does not alter the invariant splice-junction dinucleotides (GT/AG), making disruption of canonical splicing unlikely to explain the reduced mRNA levels observed. However, we cannot exclude effects on cryptic splicing^69^, and post-transcriptional mechanisms affecting mRNA stability and/or translational efficiency may contribute to reduced Lrrk2 T1348N expression. Changes in nucleotide sequence can alter local mRNA folding, thereby exposing or occluding binding sites for microRNAs and RNA-binding proteins and can influence ribosome transit rates, which in turn feeds back on mRNA stability and translational efficiency^70^. Such effects of coding nucleotide changes on mRNA structure, regulatory access and translation can therefore influence gene expression independently of changes in protein stability^71^. Supporting this notion, previous studies linked LRRK2 mRNA stability to post-transcriptional regulatory mechanisms, including microRNA-dependent control of transcript abundance^72^.

Functionally, Lrrk2 T1348N is catalytically inactive in cells as it fails to recruit/phosphorylate Rab8A and Rab10 even following induction of lysosomal damage, a well-established mechanism of LRRK2 activation^15,33,73^. Since the GTPase activity of LRRK2 regulates its association to membranes and, consequently, access to Rab substrates^74^, loss of GTP/GDP binding would be expected to impair lysosomal stress responses and autophagy-lysosome pathways^75,76^. Indeed, AP-MS of 3xFlag-tagged LRRK2 WT and T1348N ectopically expressed in HEK-293T cells revealed extensive remodeling of the LRRK2 interactome. This included loss of 14-3-3 binding, consistent with dephosphorylation at Ser910 and Ser935 (Fig. 3 and ^44,47^), and increased interactions with molecular chaperones, indicative of a less stable conformational state. Importantly, chaperone engagement may buffer conformational instability and help maintain a soluble pool of the mutant protein^48,50,77^, consistent with the comparable degradation rate and the absence of a major re-localization to insoluble fractions in our assays.

Guanosine nucleotide-binding loss results in the accumulation of the autophagic cargo p62 in macrophages and in kidney tissue in an age-dependent manner, but not in the brain or lungs (Fig. 4). This highlights that LRRK2-dependent regulation of autophagy is context-dependent and may rely on cell-type-specific interaction networks or compensatory mechanisms^78^. Of note, a similar increase in p62 levels was previously described in H4 astroglioma cells treated with the type I LRRK2 inhibitor 1 (IN-1)^79^. As type I kinase inhibitors stabilize LRRK2 in a closed active-like conformation which enhances its association with microtubules^80,81^, it is intriguing to speculate that the T1348N mutant may adopt a similar structural state. The kidney-selective phenotype observed in this study is consistent with previous reports showing that LRRK2 genetic perturbation preferentially affects kidney homeostasis, including lysosomal and autophagy-related alterations^64–66^. The stronger p62 accumulation detected in female T1348N kidneys further suggests that sex may influence the severity of tissue-specific autophagy-lysosome phenotypes, consistent with broader evidence for sex-related differences in the autophagic response to stressors as well as in PD and LRRK2-associated PD, although the mechanisms underlying this bias remain to be defined^82–84^.

Although previous studies reported that p62 interacts with LRRK2 and promotes its degradation via the autophagy-lysosome pathway^85,86^, p62 was not detected among confidently quantified interactors in our AP-MS datasets and its accumulation was not associated with increased lysosomal degradation of Lrrk2 (Fig. 2). Together, these findings suggest that Lrrk2 T1348N impairs autophagy through a mechanism independent of detectable p62-LRRK2 interaction in our AP-MS conditions. Mechanistically, we found that p62 accumulation correlates with increased binding of LRRK2 T1348N to the E3 ubiquitin ligase TRIM32, a positive regulator of autophagy acting upstream of p62^52–54^. Lrrk2 T1348N-expressing macrophages exhibit impaired autophagy, supporting a model in which loss of catalytic activity together with altered scaffolding/interactions perturbs lysosome-dependent cargo handling. These findings are interesting in light of a study reporting that LRRK2 binds the E3 ubiquitin ligase TRIM1, which competes with 14-3-3 proteins for LRRK2 binding^87^. By analogy, if a similar competitive mechanism operates between LRRK2 and TRIM32, the loss of 14-3-3 binding in the T1348N mutant could enhance TRIM32 association, consistent with our AP-MS observations. In addition, a functional interaction between LRRK2 and TRIM32 in the regulation of miRNA activity was previously observed^88^. This supports the idea that nucleotide binding and hydrolysis governs the composition of the LRRK2 interactome^89^ and consequently, signaling properties.

Although several of the observed phenotypes might initially be interpreted as loss of LRRK2 function, a key insight from our study is that genetic disruption of GTP/GDP binding does not phenocopy LRRK2 KO. In contrast to LRRK2-deficient systems, where p62 levels are reduced and autophagic flux is increased as shown here and elsewhere (Fig. 4C and ^14,90^), the T1348N mutant shows p62 accumulation and delayed flux. This divergence suggests that the presence of a structurally intact but functionally compromised LRRK2 protein exerts dominant-negative effects, likely mediated by its scaffolding domains and altered protein-protein interactions.

This distinction is particularly relevant in the context of LRRK2-targeted therapies, including small-molecule kinase inhibitors and antisense oligonucleotides (ASO) approaches, which aim to reduce LRRK2 activity or abundance in patients^10^. While reduced LRRK2 protein levels in humans are not associated with overt pathology^91^, pharmacological inhibition of LRRK2 with type I inhibitors induces dephosphorylation at Ser910/Ser935 and loss of 14-3-3 binding, a state that is not recapitulated by genetic kinase-dead mutants such as D2017A, where Ser935 phosphorylation and 14-3-3 interactions are preserved^92^. In contrast, genetic disruption of GTP/GDP binding closely mirrors inhibitor-induced dephosphorylation, positioning the T1348N KI model as a useful and interesting system to study the long-term consequences of combined kinase inhibition and scaffold dysfunction.

These findings are also relevant to human genetics, as a naturally occurring variant at the same residue (T1348P) has been identified in human populations and has been shown to disrupt GTP binding by LRRK2 and RAB10 cellular phosphorylation in a manner similar to the T1348N substitution^33^. Taken together, our results highlight nucleotide binding as a central regulatory node controlling LRRK2 expression, signaling, and interaction networks. They further suggest that chronic alteration of LRRK2 scaffolding functions, whether genetic or pharmacological, may have tissue-specific effects on autophagy and cellular homeostasis. These findings underscore the importance of carefully distinguishing between loss of LRRK2 activity and altered LRRK2 architecture when evaluating therapeutic strategies for PD.

## EXPERIMENTAL MODEL AND SUBJECT DETAILS

### Mice

Tissue samples and primary BMDMs were obtained from C57BL/6NJ WT and Lrrk2 T1348N KI mice (Dr Dave Kuldip and The Jackson Laboratory, strain #021829 [Lrrk2tm2.1Mjff]). Mice were housed under standard conditions (12 h light/dark cycle), with food and water ad libitum. All animal procedures were performed in compliance with national and institutional guidelines and were approved by the Ethical Committee of the University of Padova and the Italian Ministry of Health (License D2784.N.DMJ).

Sex and age for each experiment are reported in the corresponding figure legends. For tissue collection, animals were euthanized by cervical dislocation. Brain, lungs and kidneys were dissected; one half of each tissue was snap-frozen in liquid nitrogen and processed for Western Blotting (WB), and the remaining half was fixed for histological processing. The protocol for BMDM isolation and differentiation is described below.

### Cell lines

RAW 264.7 WT (SC-6003), Lrrk2 KO (SC-6004), and Lrrk2 T1348N KI (SC-6005) as well as HEK-293T cells were purchased from ATCC. Cells were maintained at 37°C in a humidified 5% CO_2_ atmosphere in Dulbecco’s modified Eagle’s medium (DMEM, Life Technologies) supplemented with 10% heat-inactivated fetal bovine serum (FBS, Life Technologies) and 1% penicillin/streptomycin (P/S, Life Technologies). 0.25% trypsin (Life Technologies), supplemented with 0.53 mM EDTA, was employed to generate subcultures.

## METHODS DETAILS

### Plasmids and reagents

pCHMWS-3xFlag LRRK2 WT and LRRK2 T1348N vectors were previously described^93^. The tandem fluorescent autophagy reporter mCherry-GFP-LC3 by Prof. Terje Johansen^94^ was a kind gift of Dr Sabine Hilfiker.

Pharmacological treatments were as follows: chloroquine (CQ, 50 µM, 2 or 16 h, as indicated in figure legends; Sigma-Aldrich, C6628), MG132 (20 µM, 16 h; SCBT, sc-201270), MLi-2 (0.1 μM, 90 min; Tocris Bioscience, 5756), L-leucyl-L-leucine methyl ester (LLOMe, 1 mM, 30 min; Bachem Products^16^), cycloheximide (CHX, 1 ng/μL, time points as indicated; SCBT, sc-3508).

### Murine BMDM isolation and differentiation

BMDMs were obtained from mice adapting the protocol described by Herbst and Gutierrez^95^. Briefly, bone marrow was isolated from the hind legs of 8/10-week-old mice and subjected to red blood cell lysis using red blood cell lysis buffer (Hybri-Max, Sigma-Aldrich, R7757) for 10 min at room temperature (RT). Cells were plated in differentiation medium consisting of RPMI 1640 (Life Technologies) supplemented with 2 mM GlutaMAX (Life Technologies), 25 mM HEPES (ThermoFisher Scientific), 10% heat-inactivated FBS and recombinant mouse Granulocyte-Macrophage Colony-Stimulating Factor (GM-CSF, 50 ng/mL; ThermoFisher Scientific, PMC2015), and cultured at 37°C in a humidified 5% CO_2_ atmosphere for 7 days, with medium replacement at day 3. On day 7, cells were washed, detached in ice-cold Phosphate-Buffered Saline (PBS) without Mg^2+^/Ca^2+^ (ThermoFisher Scientific, 10010023), and replated in RPMI 1640 containing 10% heat-inactivated FBS for downstream experiments.

### Immortalized cell line transfection

For AP-MS experiments, HEK-293T cells were transiently transfected using polyethyleneimine (PEI, Polysciences) at a DNA:PEI ratio of 1:2. Cells were plated on 150-mm dishes and transfected with 20 μg of 3xFlag-LRRK2 WT or 40 μg of 3xFlag-LRRK2 T1348N plasmid DNA, as determined by preliminary titration to achieve comparable bait recovery in anti-Flag immunoprecipitates. At 48-72 h post-transfection, cells were subjected to treatment with vehicle, CQ (as previously described) or starvation (16 h serum deprivation followed by 2 h in Hanks’ Balanced Salt Solution, HBSS, Life Technologies) prior to lysis and anti-Flag affinity purification for MS or immunoblot validation.

For autophagic flux measurements, RAW 264.7 WT, Lrrk2 KO and Lrrk2 T1348N cells were cultured onto 12mm glass coverslips in 24-well plates that had previously been coated with poly-L-lysine (Sigma-Aldrich), and transiently transfected with 1 μg mCherry-GFP-LC3 plasmid DNA per well using Lipofectamine 2000 (Invitrogen) at a DNA:Lipofectamine ratio of 1:2. Cells were treated with vehicle or CQ (as previously described) prior to fixation in 4% paraformaldehyde (PFA, Sigma-Aldrich) in PBS (pH 7.4) for 20 min at RT, mounting and acquisition on a Zeiss LSM700 laser scanning confocal microscope.

### Protein immunoprecipitation for AP-MS

HEK-293T cells were harvested 48-72 h post-transfection and lysed in 20 mM Tris-HCl (pH 7.5), 150 mM NaCl, 1 mM EDTA, 2.5 mM sodium pyrophosphate, 1 mM β-glycerophosphate, 1 mM sodium orthovanadate and 1% Tween 20, supplemented with protease inhibitor cocktail (Sigma-Aldrich) and phosphatase inhibitor cocktail (Life Technologies). Lysates were incubated on ice for 30 min and clarified by centrifugation at 20,000 × g for 30 min at 4°C. Supernatants were incubated for 2 h at 4°C with anti-Flag^®^ M2 affinity gel (40 μL slurry per sample; Sigma-Aldrich). Immunocomplexes were washed 10 times using buffers of decreasing ionic strength and processed either for immunoblotting or for LC-MS/MS.

### Solubility assay

To separate Triton X-100 soluble and insoluble fractions, RAW 264.7 WT and Lrrk2 T1348N cells were grown in 6-well plates and lysed in 20 mM Tris-HCl (pH 7.5), 150 mM NaCl, 1 mM EDTA, 2.5 mM sodium pyrophosphate, 1 mM β-glycerophosphate, 1 mM sodium orthovanadate and 1% Triton X-100, supplemented with protease and phosphatase inhibitor cocktails. Lysates were incubated on ice for 30 min and clarified by centrifugation at 20,000 × g for 30 min at 4°C. The supernatant was collected as the Triton-soluble fraction while the pellet constituted the insoluble fraction. Pellets were resuspended in lysis buffer containing 2% Sodium Dodecyl Sulphate (SDS) and incubated at RT for 15 min. Debris were removed by centrifugation at 20,000 × g for 30 min at 4°C. Both fractions were analyzed by SDS-PAGE and immunoblotting.

### Western Blot

RAW 264.7 cells, BMDMs and mouse tissues were lysed in RIPA buffer (20 mM Tris-HCl pH 7.5, 150 mM NaCl, 1 mM EDTA, 1 mM EGTA, 1% Nonidet^TM^ P40, 1% sodium deoxycholate) supplemented with protease inhibitor cocktail (Sigma-Aldrich) and phosphatase inhibitor cocktail (Life Technologies). Lysates were clarified by centrifugation at 20,000 × g for 30 min at 4°C. Protein concentration was determined using the Pierce^®^ BCA Protein Assay Kit (Thermo Scientific). Samples were resolved on 4-20% or 8-16% Tris-MOPS-SDS gels (GenScript) and transferred to polyvinylidene fluoride (PVDF) membranes using a semi-dry Trans-Blot^®^ Turbo^TM^ System (Bio-Rad). Membranes were blocked in 5% non-fat milk in Tris-buffered saline plus 0.1% Tween (TBS-T) for 1 h at RT under agitation and subsequently incubated with primary antibodies for 2 h at RT or overnight at 4°C. After three washes in TBS-T, membranes were incubated with horse-radish peroxidase (HRP)-conjugated secondary antibodies for 1 h at RT, and washed three additional times prior to acquisition. Immunoreactive proteins were visualized using Immobilon^®^ Classico Western HRP Substrate (Millipore) or Immobilon^®^ Forte Western HRP Substrate (Millipore) by the Imager CHEMI Premium detector (VWR). Images were acquired in .tiff format and the densitometric analysis of the detected bands was performed using the Fiji software. The following antibodies were used: rabbit anti-Lrrk2 (ab133475, abcam, 1:300 to 1:1,000), rabbit anti-LC3B (NB100-2220, Novus Biologicals, 1:5,000), rabbit anti-p62 (ab109012, abcam, 1:5,000), rabbit anti-phospho-Ser^935^–Lrrk2 (ab133450, abcam, 1:500), rabbit anti-phospho-Thr^73^-Rab10 (ab230261, abcam, 1:300), rabbit anti-14-3-3γ (PA5-29690, Invitrogen, 1:2,000), mouse anti-14-3-3pan (sc-133233, SCBT, 1:3,000), mouse anti-β-actin (A1978, Sigma-Aldrich, 1:10,000), rabbit anti-β-tubulin III (T8578, Sigma-Aldrich, 1:5,000), anti-Flag M2-Peroxidase (A8592, 1:50,000), mouse anti-Hsp70 (H5147, Sigma-Aldrich, 1:5,000), mouse anti-Hsp90 (610419, BD Biosciences, 1:2,000), rabbit anti-TRIM32 (10326-1-AP, Proteintech, 1:500); goat anti-mouse IgG-HRP (A9044, Sigma-Aldrich, 1:80,000), goat anti-rabbit IgG-HRP (A9169, Sigma-Aldrich, 1:16,000).

### RNA extraction and Real Time-PCR

Five replicates per genotype (Lrrk2 WT, KO and T1348N) were grown in 10-cm dishes to ∼ 80% confluency, washed twice with Dulbecco’s PBS (DPBS), and lysed in QIAzol^®^. Total RNA was extracted from RAW 264.7 cells using the RNeasy kit (Qiagen) according to the manufacturer’s instructions. RNA concentration and purity were assessed by a NanoDrop ND-1000 Spectrophotometer V3.3.0 (A260/280 and A260/230), and RNA integrity was verified by bleach agarose gel electrophoresis^96^. cDNA was synthesized via the ImProm-II™ Reverse Transcription System (Promega). *Lrrk2* mRNA expression levels were measured by Real-Time PCR (RT-PCR) in a CFX96 Touch-Real Time PCR Detection System (Bio-Rad). Ten ng of cDNA were amplified using the PowerUp™ SYBR™ Green Master Mix (ThermoFisher Scientific). The amplification protocol consisted of 50 °C for 2 min, 95 °C for 2 min, followed by 40 cycles at 95 °C for 15 s and 60 °C for 1 min. *Lrrk2* primers were designed within the N-terminus of the LRR domain (i.e., the epitope region recognized by the MJFF2 antibody), and *Tfrc* was used as a housekeeping gene. The following forward (Fw) and reverse (Rv) primer sequences were utilized:

*Lrrk2* Fw, AAGTCCAACTCAATTAGTGTAGGGGAAGT;

*Lrrk2* Rv, AGAACACATCACGTCGTTAGACCTATCT;

*Tfrc* Fw, TATAAGCTTTGGGTGGGAGGCA;

*Tfrc* Rv, TCATACACCCGGTTTAGCCTTGCT.

Relative expression analysis was performed using REST (http://rest.gene-quantification.info/).

### Lysosomal damage-induced recruitment assays

RAW 264.7 cells, seeded in tissue culture-treated PhenoPlates (6007710, revvity) were treated with LLOMe as previously described. The cells were fixed in 4% methanol-free PFA (15710, Electron Microscopy Sciences) in PBS for 15 min at 4°C and processed for immunofluorescence. To stain for Rab recruitment, the samples were permeabilized and blocked with 0.3% Triton X-100, 5% FCS in PBS for 20 min. To stain for Lrrk2 recruitment, the samples were permeabilized in ice-cold methanol for 10 min, followed by blocking in 5 % FCS in PBS for 20 min. Primary antibodies were diluted in PBS containing 5% FCS and incubated for 1 h at RT. The samples were washed three times in PBS, and incubated for 45 min at RT with secondary antibodies diluted in 5% FCS in PBS. After three more washes with PBS, nuclear staining was performed using 300 nM DAPI (Life Technologies, D3571) in PBS for 10 min. Images were acquired in PBS on an OPERA Phenix high-content imaging system. The images were analyzed for the number of spots per cell using Harmony High-Content Imaging and Analysis Software (revvity). The following antibodies were used: rabbit-anti-Rab10 (8127, Cell Signaling Technology), rabbit-anti-Rab8A (6975, Cell Signaling Technology), and rabbit-anti-LRRK2 (ab133474, Abcam).

### Immunofluorescence & Immunohistochemistry

Mouse tissues were fixed using 4% PFA (Sigma-Aldrich) in PBS (pH 7.4) for 48 h at 4°C. Tissues were cryoprotected in 30% sucrose in PBS, embedded in OCT (Optimal Cutting Temperature, Kaltek), and cryo-sectioned. Kidneys were sectioned at 6 μm and collected on Superfrost Plus slides (ThermoFisher Scientific). Sections were incubated for 10 min in a quenching solution (50 mM NH_4_Cl in PBS), rinsed three times in PBS, and treated with Sudan Black for 15 min to reduce tissue autofluorescence. After three additional washes in PBS, slices were saturated for 1 h at RT in a blocking buffer (2% BSA, 15% goat serum, 0.25% gelatine, 0.20% glycine and 0.5% Triton X-100) and incubated overnight at 4°C with primary antibodies diluted in blocking solution. The following day, sections were washed three times in PBS and incubated with secondary antibodies for 1 h at RT, counterstained with Hoechst-33258 (1:10,000 in PBS; Invitrogen) for 5 min, and mounted with Mowiol (Calbiochem). Z-stack images (z-stack thickness, namely z-step × n slices, 4 µm) were obtained on Zeiss LSM700 laser scanning confocal microscope.

The following antibodies were used: rabbit anti-p62 (ab109012, abcam, 1:200), rabbit anti-TRIM32 (10326-1-AP, Proteintech, 1:200), rat anti-LAMP1 (ab25245, abcam, 1:200); goat anti-rabbit Alexa Fluor 488 (A11034, Invitrogen, 1:200), goat anti-rat Alexa Fluor 647 (A21247, Invitrogen, 1:200).

### Protein digestion and LC-MS/MS analysis

Samples from immunoprecipitation experiments were eluted by resuspending beads in 4× Laemmli sample buffer (200 mM Tris-HCl pH 6.8, 8% SDS, 400 mM DTT, 40% glycerol, bromophenol blue), centrifuged at 7,500 × g for 15 min to pellet the beads, and eluates were resolved by SDS-PAGE on 4-20% Tris-MOPS precast gels (GenScript) for electrophoretic separation. Samples were run for about 1 cm in length, and lanes were excised and split into two fractions for protein digestion. Gel slices were cut into small pieces and proteins were reduced with dithiothreitol (10 mM DTT in 50 mM NH4HCO3, for 1 h at 56 °C), alkylated with iodoacetamide (55 mM IAA in 50 mM NH4HCO3, for 45 min at RT and in the dark), and digested with sequencing grade modified trypsin (Promega, 12.5 ng/μL in 50 mM NH4HCO3). Peptides were eluted from the gel using 50% acetonitrile (ACN)/0.1% formic acid (FA) and vacuum-dried in a speed-vac. Samples were resuspended in 30 µL of 3% ACN/0.1% FA, and 6 µL were analyzed with a LTQ-Orbitrap XL mass spectrometer (Thermo Fisher Scientific) coupled to a HPLC UltiMate 3000 (Dionex – Thermo Fisher Scientific) through a nanospray interface. Peptides were separated in an 11-cm-long capillary column (PicoFrit, 75-μm ID, 15-μm tip, New Objective) packed in-house with C18 material (Aeris Peptide 3.6 μm XB C18, Phenomenex) using a linear gradient of ACN/0.1% FA from 3 to 40% in 40 min at a flow rate of 250 nL/min. The instrument operated in a data-dependent acquisition mode with a Top10 method (one full MS scan at 60,000 resolution in the Orbitrap, followed by the acquisition of the MS/MS spectra of the ten most intense ions in the linear ion trap). Two technical replicates were acquired for each biological replicate.

Raw data files were processed using MaxQuant software v1.5.1.2 with Andromeda search engine^97^. Proteins were searched against the human section of the UniProt database (UP000005640, version 2020-09-30, 75074 entries), concatenated with a database of common contaminants found in proteomic experiments. Trypsin was selected as enzyme with up to two missed cleavages allowed, and cysteine carbamidomethylation and methionine oxidation were set as fixed and variable modifications, respectively. A minimum of two peptides was required for protein identification, and results were filtered considering a false-discovery rate (FDR) of 0.01, both at the peptide and protein levels.

### Differential abundance analysis

For proteins to be considered for downstream analysis, the detection criteria were ≥2 unique peptides across the MS dataset and ≥2 peptides across technical replicates for the corresponding assigned protein. This resulted in 188 proteins, including LRRK2. Intensity data were mean averaged across technical replicates.

The DEP R package^98^ was used to determine differential abundance of proteins between experimental conditions. Data preprocessing included removal of proteins which were not represented across biological replicates (145 proteins retained), normalization using variance stabilizing transformation and imputation using Bayesian principal component analysis. Pairwise comparisons were performed across LRRK2 genotype (wild type vs T1348N mutant) using linear models based on the limma R package^99^ to determine differential abundance.

### Statistical analysis

All quantitative data are presented as mean ± SEM (standard error of the mean). Statistical analyses were performed using GraphPad Prism v10.6.1. The number of technical or biological replicates (*n*) and the statistical tests used for each experiment are reported in the corresponding figure legends. Unless otherwise stated, all tests were two-tailed. Prior to statistical testing, datasets were assessed for normality and screened for outliers. For comparisons between two groups, an unpaired two-tailed Student’s t test was used. For comparisons among three or more groups, one-way ANOVA followed by Tukey’s multiple-comparisons test was used. For two-factor analyses (e.g., genotype × age), two-way ANOVA followed by Šídák’s multiple-comparisons test was used. When assumptions of parametric testing were not met, non-parametric tests were applied (Kruskal-Wallis test with Dunn’s multiple-comparisons test). For datasets expressed relative to a reference condition set to 1 (e.g., WT-normalized fold change), a two-tailed one-sample t test versus the hypothetical value of 1 was used when appropriate. For immunofluorescence quantifications, multiple fields were quantified per mouse and averaged to generate a single value per mouse (details and *n* reported in figure legends). For the tandem mCherry-GFP-LC3 autophagic flux assay, image acquisition and analysis were performed blinded; z-stacks were acquired and quantified in Fiji following maximum-intensity projection, background subtraction/thresholding, and automated puncta detection/overlap quantification using ComDet v0.4.2. Significance thresholds are denoted as ns, not significant; * p < 0.05, ** p < 0.01, *** p < 0.001, **** p < 0.0001.

## Acknowledgements

This work was funded by a grant from the Michael J. Fox Foundation for Parkinson’s research (grant number 18285) to PAL and EG, and was supported by the European Union – NextGenerationEU (NRRP, Mission 4, Component 2, Investment 1.1 – PRIN, Project 20222LRHCW, CUP C53D23005560006) to EG and by a CARIPARO PhD Fellowship and an Aldo Gini Fellowship to SC. This work was additionally supported by the Francis Crick Institute (to MGG), which receives its core funding from Cancer Research UK (CC2081), the UK Medical Research Council (CC2081), and the Wellcome Trust (CC2081).

## CRediT author statement

Conceptualization, S.H., P.A.L., E.G., and S.C.; Methodology, G.F., S.H., A.M., I.B., J.E.T., D.T., and S.C.; Investigation, G.F., S.H., A.M., G.T., L.I., I.B., N.P., and S.C.; Formal analysis, G.F., S.H., A.M., I.B., J.E.T., and S.C.; Data curation, G.F., S.H., A.M., I.B., J.E.T., and S.C.; Resources, M.G. G., G.A., C.M., P.A.L., and E.G.; Visualization, G.F., S.H., A.M., J.E.T., and S.C.; Writing – original draft, G.F., P.A.L., E.G, and S.C.; Writing – review & editing, all authors; Supervision, P.A.L., E.G., and S.C.; Project administration, P.A.L., E.G. and S.C.; Funding acquisition, P.A.L., and E.G.

---

**Supplementary Figure 1.**
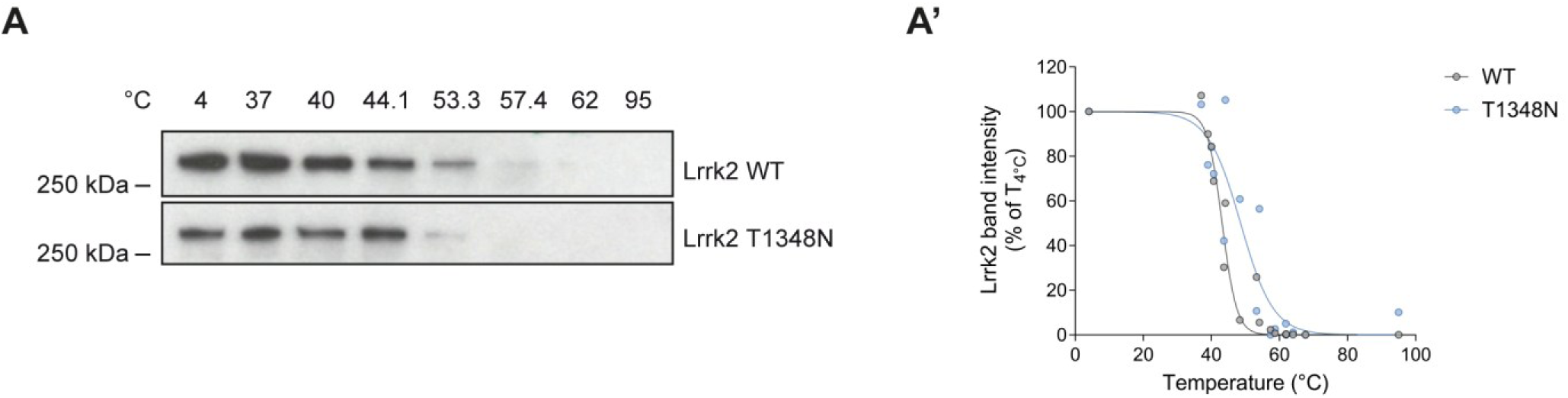
CETSA indicates no consistent shift in apparent thermal stability of Lrrk2 T1348N. (**A**) Cellular thermal shift assay (CETSA)/thermal denaturation profiling of endogenous Lrrk2 in RAW 264.7 WT and T1348N cells across the indicated temperature range, assessed by immunoblotting. (**A’**) Quantification of Lrrk2 band intensity from (**A**), expressed as percentage of the 4°C condition and plotted as a function of temperature. Data are presented as individual measurements and are representative of two independent experiments; curves are shown as fitted trends.

**Supplementary Figure 2.**
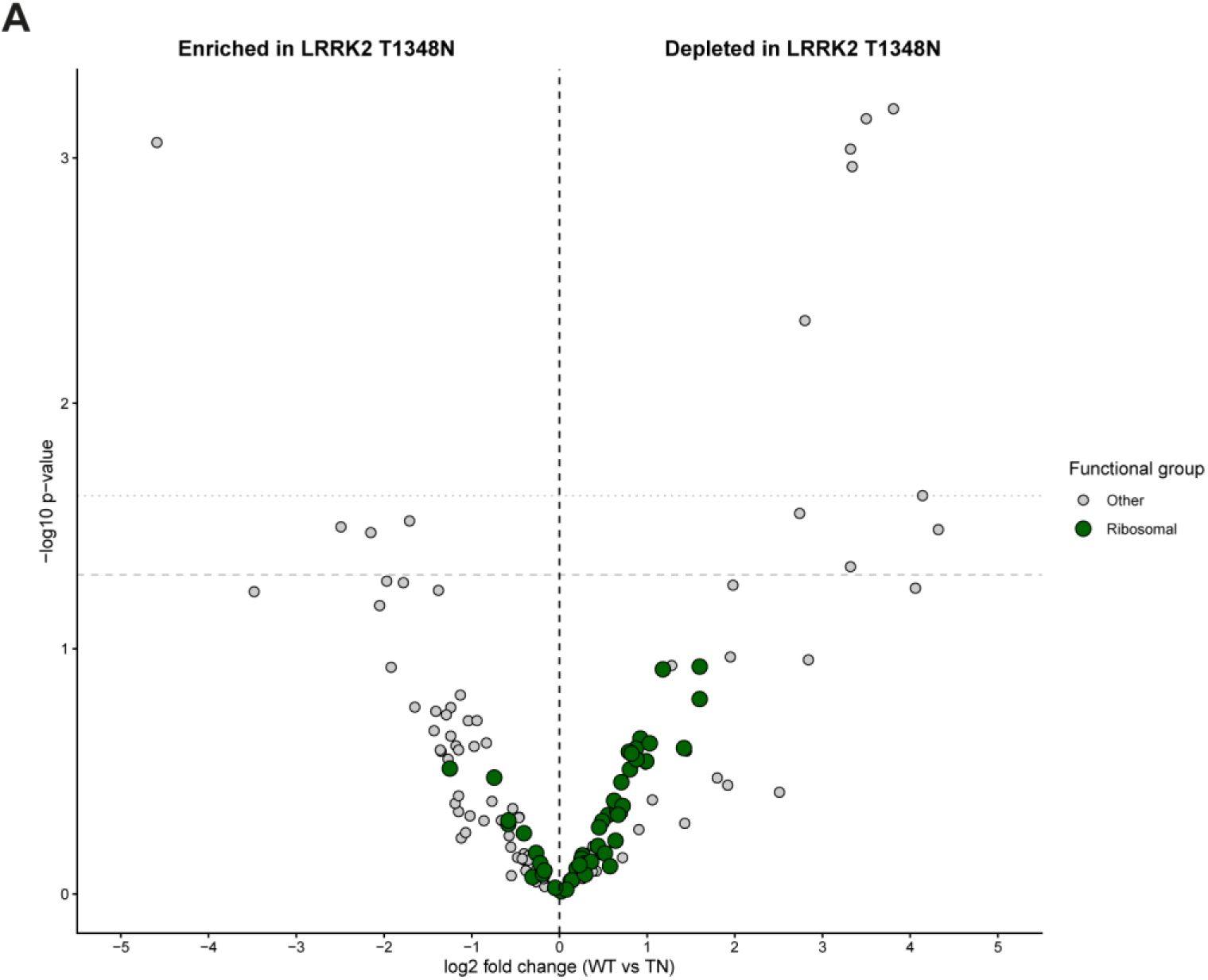
Ribosomal proteins show heterogeneous changes in the LRRK2 T1348N interactome. Volcano plot showing differential enrichment of proteins co-purified with 3xFlag-LRRK2 WT or T1348N under basal conditions, with ribosomal proteins highlighted. Data are plotted as log2 fold-change (WT/T1348N) against -log10 p-value from *n* = 2 biological replicates per genotype; negative values indicate enrichment with LRRK2 T1348N, whereas positive values indicate reduced association with LRRK2 T1348N. Ribosomal proteins are distributed on both sides of the plot, consistent with heterogeneous changes rather than a uniform shift in association with either WT or T1348N LRRK2. Each dot represents one quantified protein.

